# Impact of peripheral mutations on the access channels of human cytochrome P450 1A2

**DOI:** 10.1101/754739

**Authors:** Beili Ying, Yang Zhong, Jingfang Wang

## Abstract

As an important member of cytochrome P450 (CYP) enzymes, human CYP1A2 is associated with the metabolism of caffeine and melatonin and the activation of precarcinogens. Besides, this CYP protein also involves in metabolizing 5-10% of clinical medicines. Some peripheral mutations in CYP1A2 (P42R, I386F, R431W, and R456H) significantly decrease the enzyme activities, resulting in a vital reduction in substrate metabolisms. To explore the effects of these peripheral mutations, we constructed a membrane-binding model for the full-length human CYP1A2 and studied their dynamic behaviors on lipid membranes. Free energy calculations indicate that the peripheral mutations donot influence substrate binding. P42R is located in the N-terminal anchor, and its positive charged sidechain is adverse to membrane binding. I386F enhances the van der Waals contacts of the water channel bottleneck and R456H breaks the hydrogen bonding interactions that function to position the BC loop, both of which result in a significant inhibition on the water channel. R431W causes a sidechain conformational rearrangement for aromatic residues around the substrate channel, making it in a closed state in most cases. Our computational simulations demonstrate that pi-pi stacking interactions are essential for substrate binding and channel opening. We hope that these findings may be of general relevance to the mutation-induced activity changes for CYP proteins, providing useful information for understanding the CYP-mediated drug metabolism.

## Introduction

Cytochrome P450 (CYP) enzymes are a large family of heme-containing proteins that are widely distributed in all kingdoms of life with responsibility for metabolizing a wide range of structurally dissimilar substrates (Danielson, 2002). In mammal cells, CYP proteins are usually located on the cell membranes of the endoplasmic reticulum, acting as mono-oxygenases to add or unmask a polar group into substrates, making them more soluble in water for further metabolisms and clearances (Chen et al., 2011). In most human cells, CYP proteins are major enzymes for phase I drug metabolisms, accounting for more than 90% clinically used medicines (Wang et al., 2009). In addition to drug metabolism, CYP proteins are also essential for controlling the levels of cholesterol, steroids, as well as lipids (for example, prostacyclins and thromboxane A2). Because these proteins are related to vascular autoregulation and the formation of cholesterol, steroids, and arachidonic acid metabolites (Ingelman-Sundberg, 2005). Of note, CYP proteins are also well-known for their single nucleotide polymorphisms (SNP), which is estimated to have a significant effect on more than 30% of clinical therapies (Ingelman-Sundberg, 2004; Shou, 2004).

As an important member of CYP1 family, human CYP1A2 is predominantly expressed in liver cells, secondary in the intestine, pancreas, and brain. This CYP protein takes charge of the metabolisms of caffeine, melatonin, and nearly 5-10% of clinical medicines, including flutamide, lidocaine, as well as triamterene (Faber et al., 2005). Besides, CYP1A2 also plays a crucial role in the activation of precarcinogens, including aromatic and heterocyclic amines, and polycyclic aromatic hydrocarbons (Eaton et al., 1995). Thus, the overexpression and dysfunction of human CYP1A2 are often associated with high risks of cancers and other diseases (Bugano et al., 2008; Cornelis et al., 2006; Jernstrom et al., 2008). To date, 16 genetic polymorphisms of human CYP1A2 have been identified (Allorge et al., 2003; Chevalier et al., 2001; Huang et al., 1999; Murayama et al., 2004; Saito et al., 2005; Zhou et al., 2004), among which 8 alleles significantly decrease the enzymatic activities (**Table 1**). These genetic polymorphisms (P42R, F186L, I386F, R431W, and R456H) are peripheral mutations far away from the active pocket and have faint influences on the protein expressions.

**Table 1.** The genetic polymorphisms of human CYP1A2. The allele 1 is referred to the wild type, and the SNPs are ordered based on their discovery time.

| Allele | Nucleotide Changes | AA Mutations | Activity Changes | Ref |
| --- | --- | --- | --- | --- |
| 1 | Wild-type | — | — | — |
| 2 | 63C>A | F21L | — | 12 |
| 3 | 2385G>A, 5347T>C | D348N | Decreased | 13 |
| 4 | 2499A>T | I386F | Significant Decreased | 13 |
| 5 | 3497G>A | C406Y | — | 14 |
| 6 | 5090C>T | R431W | Significant Decreased | 13 |
| 7 | 3533G>A | — | Significant Decreased | 15 |
| 8 | 5166G>A, 5347T>C | R456H | Significant Decreased | 16 |
| 9 | 248C>T | T83M | — | 17 |
| 10 | 502G>C | E168Q | — | 17 |
| 11 | 558C>A | F186L | Significant Decreased | 17 |
| 12 | 634A>T | S212C | — | 17 |
| 13 | 1514G>A | G299S | — | 17 |
| 14 | 5112C>T | T438I | Normal | 17 |
| 15 | 125C>G, 5347T>C | P42R | Significant Decreased | 15 |
| 16 | 2473G>A, 5347T>C | R377Q | Decreased | 15 |

To explore the effect of the peripheral mutation F186L, Zhang et al. performed molecular dynamics simulations on the crystal structure of human CYP1A2 (PDB entry 2hi4) (Sansen et al., 2007). In this crystal structure, CYP1A2 was modified to remove the N-terminal transmembrane region to improve solubility and to reduce aggregation. However, the N-terminal transmembrane region is essential for recognizing and binding to the cell membranes of the endoplasmic reticulum (Jang et al., 2010; Park et al., 2015). In the current study, we remodeled the N-terminal transmembrane region and constructed a membrane-binding model for the full-length human CYP1A2. Based on this membrane-binding model, we also performed molecular dynamics simulations and free energy calculations on the wild type and mutant CYP1A2 in complex with α-nappthoflavone (ANF), intending to study their dynamic behaviors on lipid membranes.

## Materials and Computational Methods

### 1. Membrane-binding models for the wild type and mutant CYP1A2

The full-length sequence of the wild type human CYP1A2 was obtained from the UniProt database with an entry P05177 (UniProt Consortium, 2018). The catalytic domain was taken from the crystal structure of CYP1A2 (PDB entry 2hi4) in complex with ANF (Sansen et al., 2007). The N-terminal anchor region was constructed by homology modeling using Modeller 9.22 (Sali and Blundell, 1993), and subsequently inserted into dioleoyl-sn-glycero-3-phosphoethanolamine (POPC) membranes. The orientation of the wild type CYP1A2 on POPC membranes was obtained from the CYP-membrane database (http://cyp.upol.cz) (Srejber et al., 2018). The P42R, F186L, I386F, R431W, and R456H mutant structures were constructed based on the membrane-binding model of the wild type CYP1A2 using Modeller 9.22.

### 2. ANF-binding models for the wild type and mutant CYP1A2

The chemical structure of ANF was abstracted from the crystal structure of human CYP1A2 (PDB entry 2hi4) (Sansen et al., 2007). The docking files for the wild type and mutant CYP1A2 and ANF were prepared by AutoDock Tools (Morris et al., 2009). In the docking procedures, the docking grid was set up to 40, 40, and 40 points in x, y, and z directions with a grid spacing of 0.375 Å, surrounding the active pocket near the heme group. Subsequently, ANF was docked to the catalytic domain of the wild type and mutant CYP1A2 structures by AutoDock Vina (Trott and Olson, 2010). Totally 10,000 docking results were generated for each structural model of CYP1A2 and clustered using a roo-mean-square (RMS) tolerance of 2.0 Å. The favorable docking result for each structural model of CYP1A2 was finally selected as the ANF-binding model.

### 3. Molecular dynamics simulation protocols

In the beginning, all the non-polar hydrogens in the models were removed, and residue-specific pKa values were calculated by the linear Poisson-Boltzmann equation with a dielectric constant of 4.0 (Burger and Ayers, 2011). The hydrogens were then re-added based on the computational residue-specific pKa values mentioned above. Then, all the computational models were solvated in the explicit TIP3P water surroundings and subjected to energy minimization for 5000 steps by steepest descent and conjugate gradient. Molecular dynamics (MD) simulations were finally performed with periodic boundary condition, NPT ensembles (310K and 1 bar) by NAMD 2.1.2 (Phillips et al., 2005) and CHARMM36m force field parameters (Huang et al., 2017). Langevin thermostat and isotropic scaling were employed to control the simulation temperature and pressure with a time constant of 0.1 ps and 1.0 ps, respectively (Grest and Kremer, 1986). Isothermal compressibility was set to 4.5×10^-5^ per bar for solvent simulation. SHAKE algorithm was used to restrict all the chemical bonds in the simulation systems with a tolerance of 10^-6^ (Krautler and Hunenberger, 2006). The particle mesh Ewald (PME) algorithm was applied to treat electrostatic interactions with an interpolation order of 4.0 and a grid spacing of 0.12 nm (Di Pierro et al., 2015). The cut-off value for van der Waals interactions was set to 12 Å, and the time step for simulations was 2 fs.

### 4. Free energy calculations for ANF binding affinity

The ANF binding affinity was estimated by molecular mechanics Poisson-Boltzmann surface area (MM-PB/SA) and molecular mechanics Generalized-Born surface area (MM-GB/SA) methods (Wang et al., 2019). In the MM-PB/SA and MM-GB/SA methods, the binding free energy alteration was calculated as the difference between the free energies of CYP1A2, ANF, and their complex. These free energies were estimated by summing the internal energy in the gas phase, the solvation free energy, and a vibrational entropy term. Because all the simulation systems have similar entropies in the current study, and the entropy contributions are neglected to compare ANF binding affinities between the wild type and mutant CYP1A2. The internal energy in the gas phase was a standard force field energy calculated by strain energies from covalent bonds and torsion angles, non-covalent van der Waals, as well as electrostatic energies. The solvation free energy was calculated by electrostatic and non-polar energies. The former was obtained from either Poisson-Boltzmann (PB) or Generalized Born (GB) methods. The latter was estimated by molecular solvent accessible surface areas (SASAs). A total of 100 snapshots were retrieved from the last 10-ns simulation trajectories with an interval of 100 ps, and free energy calculations were performed by MMPBSA.py (Miller et al., 2012).

### 5. Data analysis and structural visualization

The access channels in the wild type and mutant CYP1A2 structures were calculated and analyzed by CAVER (Petrek et al., 2006). The starting point for searching access channels was set to 3 Å above the porphyrin ring of the heme group. The access channels were named according to Wade et al.’s nomenclature (Cojocaru et al., 2007). The water channel is considered open if the bottleneck radius is more than 1.5 Å, and the substrate channel is considered open if the bottleneck radius is more than 4.0 Å. All the protein structures in the current study were visualized By PyMol (Yuan et al., 2016).

## Results

### 1. Peripheral mutations donot influence substrate binding

According to the crystallographic studies(Sansen et al., 2007), the protein architecture of human CYP1A2 shows a typical cytochrome P450 folding, composed of 12 α-helices (designated from A to L) and 3 β-strands (designated β1 to β3) with the heme group deeply buried inside the protein core (**Figure 1**). F, G, I, and K helices, together with two short helices B’ and F’, and the heme group constitute the active pocket for CYP1A2 substrates. As shown in **Figure 1,** P42R is located in the N-terminal region, which is associated with membrane binding. F186L is a peripheral mutation located on the loop between D and E helices, nearly 28.6 Å away from the heme iron. I386F and R456H are located near the water channel, while R431W is near the substrate channel. All these mutations are not in the active pocket, having no direct interactions with substrates.

**Figure 1.**
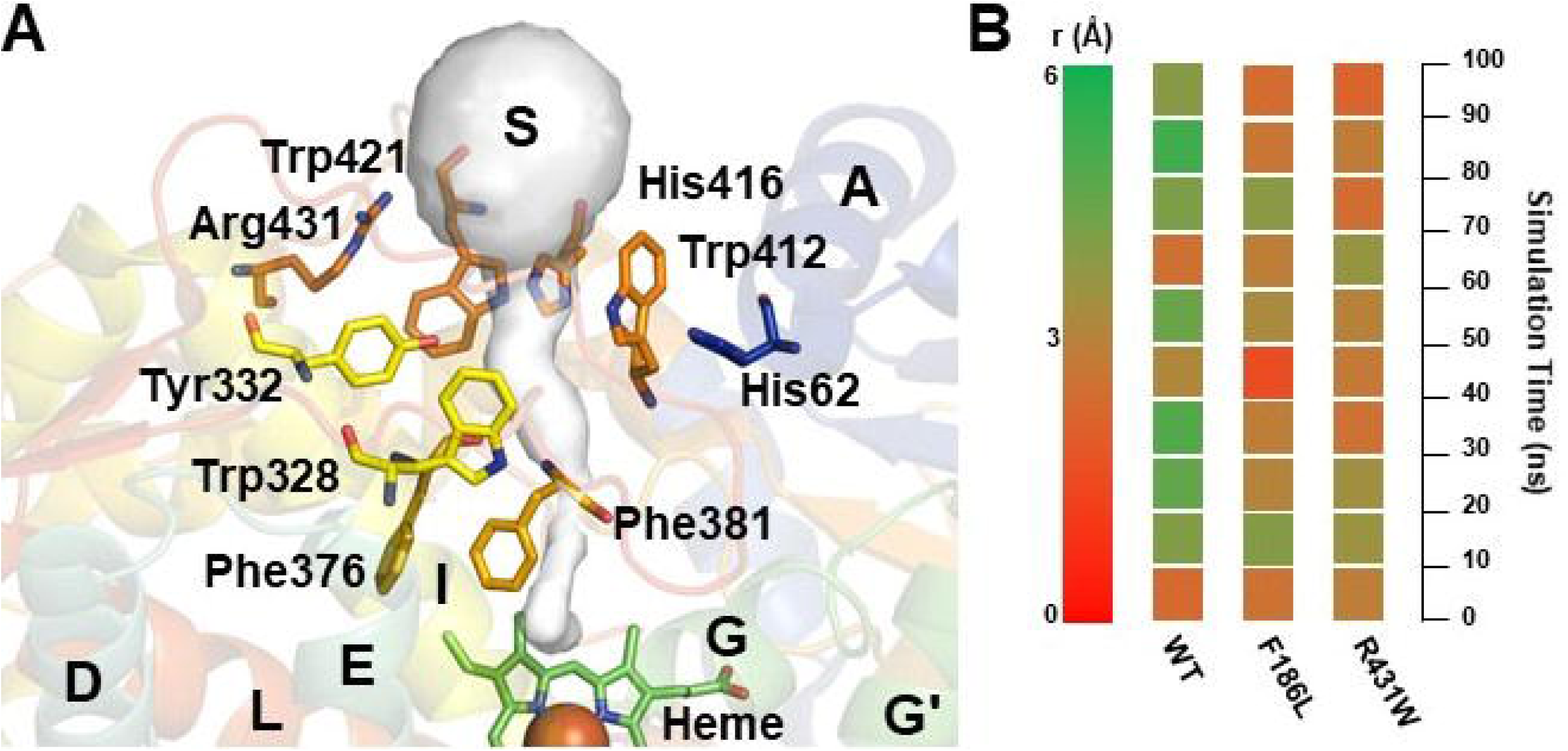
A stereo-view of the protein architecture for human CYP1A2. The protein backbone structure is shown in cartoon representations with rainbow colors. The mutation sites for P42R, F186L, I386F, R431W, and R456H are represented in sticks and are colored based on the secondary structure nearby. The substrate and water channels are shown in grey tunnels, labeled as “S” and “W” respectively.

To study the influence of these mutations on substrate binding, we also calculated the binding free energies for ANF bound to the wild type and mutant CYP1A2 by using MM-PB/SA and MM-GB/SA methods. As shown in **Table 2,** the binding free energies have no difference in statistics for ANF bound to the wild type and the mutants P42R, I386F, R431W, and R456H, indicating that these mutations (P42R, I386F, R431W, and R456H) have no impact on substrate binding. However, the binding free energy for ANF bound to F186L is significantly different from the wild type and other mutants, revealing that F186L mutation reduces the binding affinity of substrates. This is mainly because F186L mutation causes a significant reduction in the active pocket volume. In our simulations, the active pocket volume for the F186L mutant structure is about 93 ± 30 cubic angstroms, much smaller than the wild type (178 ± 29 cubic angstroms, T-test p-value < 0.05).

**Table 2.** Binding free energies (kcal/mol) for ANF bound to the wild type and mutant CYP1A2 structures.

| Energy terms | Wild type | P42R | F186L | I386F | R431W | R456H |
| --- | --- | --- | --- | --- | --- | --- |
| $\Delta E_{ele}$ | $-3.86 \pm 1.81$ | $-3.84 \pm 1.80$ | $-3.75 \pm 1.99$ | $-3.86 \pm 1.80$ | $-3.85 \pm 1.81$ | $-3.85 \pm 1.80$ |
| $\Delta E_{vdw}$ | $-15.37 \pm 1.79$ | $-15.31 \pm 1.80$ | $-13.50 \pm 1.81$ | $-15.21 \pm 1.79$ | $-15.50 \pm 1.62$ | $-15.27 \pm 1.79$ |
| $\Delta E_{gas}$ | $-19.23 \pm 3.64$ | $-19.15 \pm 3.64$ | $-19.25 \pm 3.72$ | $-19.07 \pm 3.47$ | $-19.35 \pm 3.81$ | $-19.12 \pm 3.60$ |
| $\Delta G_{non-polar/PB}$ | $-1.84 \pm 0.09$ | $-1.90 \pm 0.11$ | $-1.99 \pm 0.10$ | $-1.83 \pm 0.09$ | $-1.87 \pm 0.10$ | $-1.85 \pm 0.09$ |
| $\Delta G_{ele/PB}$ | $12.55 \pm 2.74$ | $12.02 \pm 2.69$ | $15.54 \pm 2.94$ | $12.90 \pm 2.45$ | $12.50 \pm 2.74$ | $12.63 \pm 2.62$ |
| $\Delta G_{sol/PB}$ | $10.71 \pm 2.65$ | $10.12 \pm 2.71$ | $13.55 \pm 2.98$ | $11.07 \pm 3.25$ | $10.63 \pm 2.95$ | $10.78 \pm 2.84$ |
| **$\Delta G_{bind/PB}$** | **$-8.52 \pm 3.22$** | **$-9.03 \pm 3.21$** | **$-5.70 \pm 3.07$** | **$-8.00 \pm 3.51$** | **$-8.72 \pm 3.62$** | **$-8.34 \pm 3.52$** |
| $\Delta G_{non-polar/GB}$ | $-2.48 \pm 0.10$ | $-2.50 \pm 0.10$ | $-2.65 \pm 0.15$ | $-2.50 \pm 0.10$ | $-2.50 \pm 0.10$ | $-2.48 \pm 0.10$ |
| $\Delta G_{ele/GB}$ | $11.04 \pm 1.94$ | $11.25 \pm 1.99$ | $13.97 \pm 1.96$ | $10.74 \pm 2.01$ | $11.74 \pm 1.94$ | $11.62 \pm 1.95$ |
| $\Delta G_{sol/GB}$ | $8.56 \pm 2.09$ | $8.75 \pm 2.05$ | $11.32 \pm 2.45$ | $8.24 \pm 2.00$ | $9.24 \pm 2.10$ | $9.14 \pm 2.10$ |
| **$\Delta G_{bind/GB}$** | **$-10.67 \pm 1.89$** | **$-10.40 \pm 1.90$** | **$-7.93 \pm 2.05$** | **$-10.83 \pm 2.01$** | **$-10.11 \pm 1.89$** | **$-9.98 \pm 2.27$** |

Because single mutations have little impact on the entropy of the entire protein, the entropy contributions ( $T\Delta S$ ) are neglected in the binding free energy estimations.

In the wild type, ANF well fits the size and shape of the active pocket. The aromatic residues Phe125, Phe226, Phe256, and Phe260 contribute to the binding affinity of ANF. These aromatic residues provide both orthogonal and parallel pi-pi stacking interactions with the substrate, recognizing and positioning the substrate in a favorable orientation for further metabolic reactions **(Figure 2).** In the F186L mutant, the smaller active pocket leads to a structural rearrangement of residue sidechains. Thus, the rearranged sidechains impair the pi-pi stacking interactions between ANF and the aromatic residues in the active pocket. As a result, ANF shows more flexible in the mutant structure and does not well fit the size and shape of the active pocket anymore.

**Figure 2.**
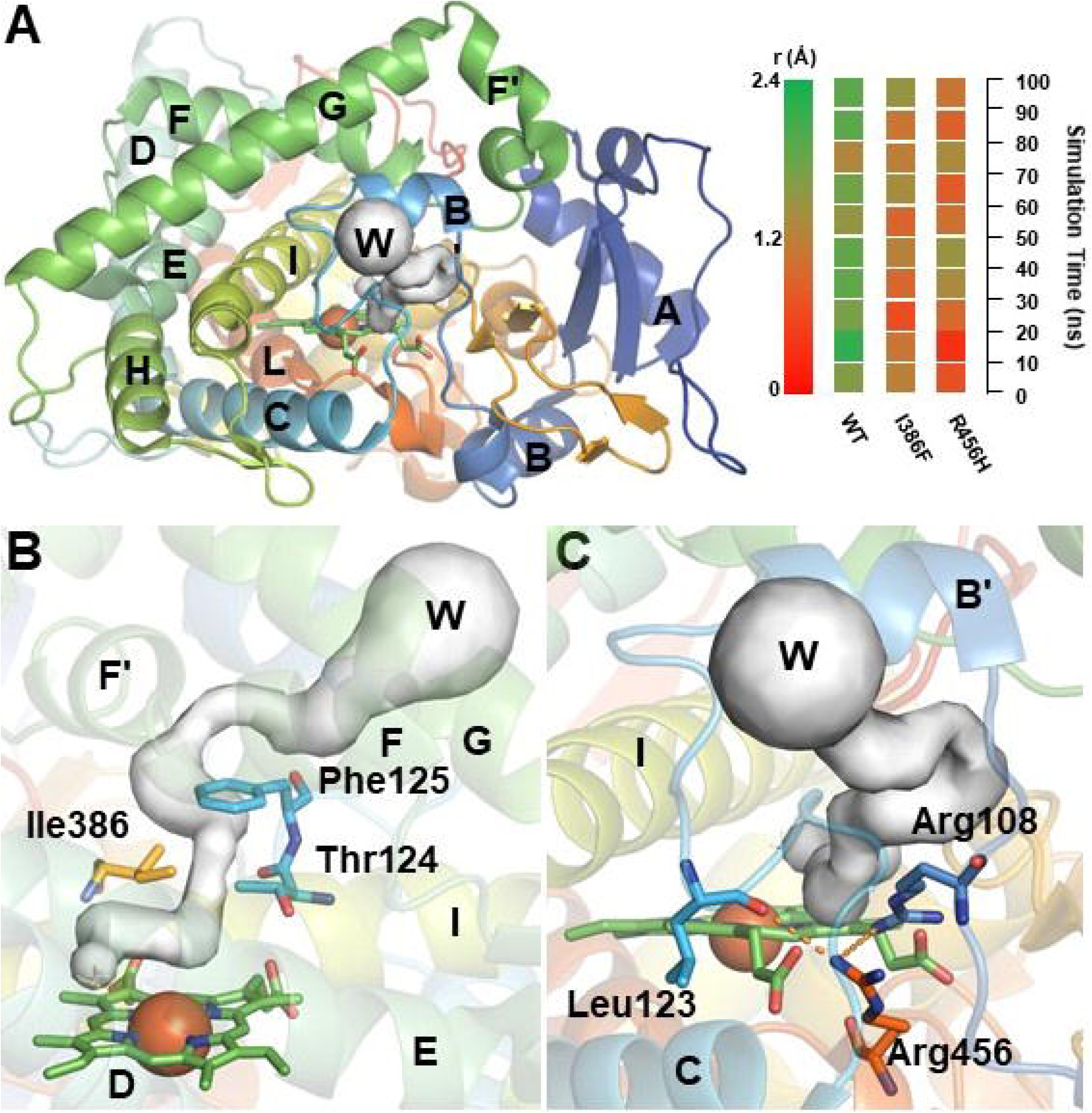
Detailed information for ANF binding in the active pocket. The protein backbone is shown in grey cartoon representations. The heme group, ANF, and the key residues are in sticks with carbon, oxygen, nitrogen, and iron colored in green, red, blue and orange, respectively. ANF is recognized and positioned in the active pocket by pi-pi stacking interactions with aromatic residues.

Additionally, Phe186 forms a pi-pi stacking interaction with Phe481 in the wild type, functioning to stabilize D and E helices and the C-terminal loop region. The C-terminal region is a key component of the substrate channel **(Figure 3)**, and the pi-pi stacking interaction between Phe186 and Phe481 has a significant impact on the substrate entering into the active pocket. The substitution from phenylalanine to leucine changes the aromatic sidechain to the aliphatic, breaking the pi-pi stacking interaction mentioned above and increasing the flexibility of D and E helices and the C-terminal loop region. Protein motion analysis indicated that the substrate channel is mostly closed in the F186L mutant, which is open in most cases in the wild type (Zhang et al., 2011).

**Figure 3.**
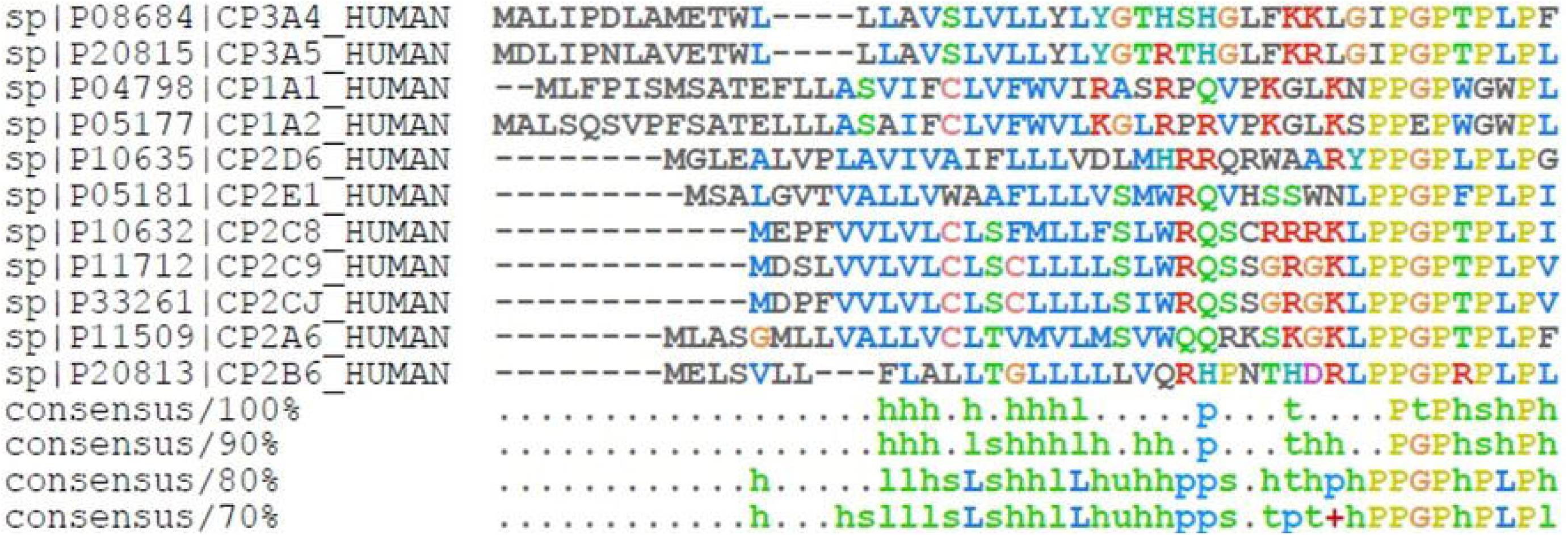
Pi-pi stacking interaction between Phe186 and Phe481 in the wild type. F186L is located on the loop between D and E helices, forming a significant pi-pi stacking interaction with Phe481 in the C-terminal loop region. This pi-pi stacking interaction functions to stabilize the C-terminal loop region, which has a significant contribution to the substrate channel (labeled as “S”).

### 2. P42R is adverse to the membrane binding of CYP1A2

Mammalian cytochrome enzymes are membrane-associated proteins, inserting in the lipid bilayers by a single N-terminal transmembrane helix (also called N-terminal anchor) (Black, 1992). Furthermore, the N-terminal anchor also contains a signal peptide that guides CYP proteins to the endoplasmic reticulum or mitochondria (Avadhani et al., 2011). Thus, the N-terminal anchor is essential for membrane binding of CYP proteins in the endoplasmic reticulum. As mentioned above, P42R is located on the N-terminal loop in the membrane anchor region. Of note, P42R is in a proline-rich motif in the N-terminal anchor **(Figure 4).** Multiple sequence alignment shows that the proline-rich motif is highly conserved in CYP1 and CYP2 families. The substitution from proline to arginine in the proline-rich motif breaks the conserved structure, resulting in instability of the N-terminal anchor. Additionally, the arginine sidechain carries positive charges and has an electrostatic exclusion effect with the phosphate group of lipid bilayers, which is adverse to the membrane binding of CYP proteins.

**Figure 4.**
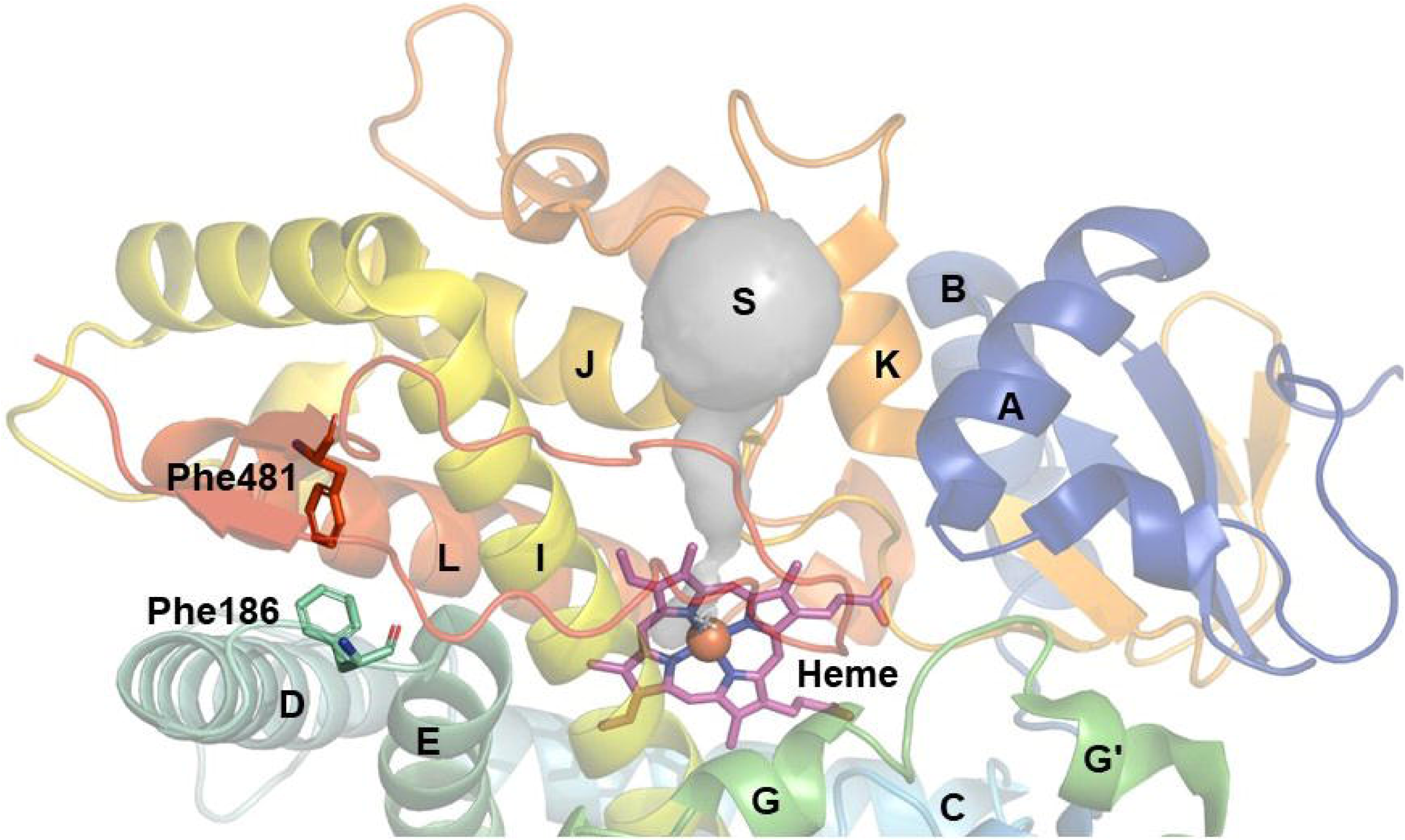
Multiple sequence alignment of the N-terminal anchor for CYP proteins. The N-terminal anchor is a transmembrane helix capable of binding to the membrane of the endoplasmic reticulum or mitochondria. The N-terminal anchor contains a proline-rich motif, which is highly conserved in most CYP proteins.

Membrane binding is essential for CYP proteins because the interactions with lipid bilayers stabilize the flexible loops and have a significant impact on the access channels for ligand exchange and water exchange between the environment and the active pocket (Berka et al., 2013; Cojocaru et al., 2011; Srejber et al., 2018). According to the previous theoretical studies, the interaction of CYP1A2 with lipid bilayers and the extended conformation of BC and FG loops facilitate the opening of the water channel (Berka et al., 2013; Cojocaru et al., 2011). Due to carrying positive charges, P42R mutation is adverse to membrane binding of CYP1A2. Thus, this mutation can influence the access channels for water and substrates, having a significant impact on both ligand and water exchanges.

### 3. I386F and R456H have a significant impact on the water channel

As essential metabolizing enzymes, CYP proteins can oxidize both xenobiotic and endogenous compounds by either adding or exposing a polar group to enhance their solubility in water for further metabolisms (Chen et al., 2011; Guengerich, 2008). The CYP-catalyzed metabolic reactions need a coenzyme NADPH (to provide hydrogen) and oxygen (to form the reactive iron-oxo intermediate Compound 1), to produce the hydroxylated metabolites (Wang et al., 2009). During the metabolic reaction, a water molecule is generated by the oxidative reaction and has to be expulsed immediately by the water channel. In the wild type CYP1A2, the water channel consists of the loop region between B and C helices (BC loop, **Figure 5A**). Small-angle X-ray scattering (SAXS) experiments and molecular dynamics simulations indicate that the BC loop is one of the membrane-interacting regions, and CYP1A2 has the most contact area for the BC loop among all the CYP proteins (Berka et al., 2013; Skar-Gislinge et al., 2015; Yang et al., 2017).

**Figure 5.**
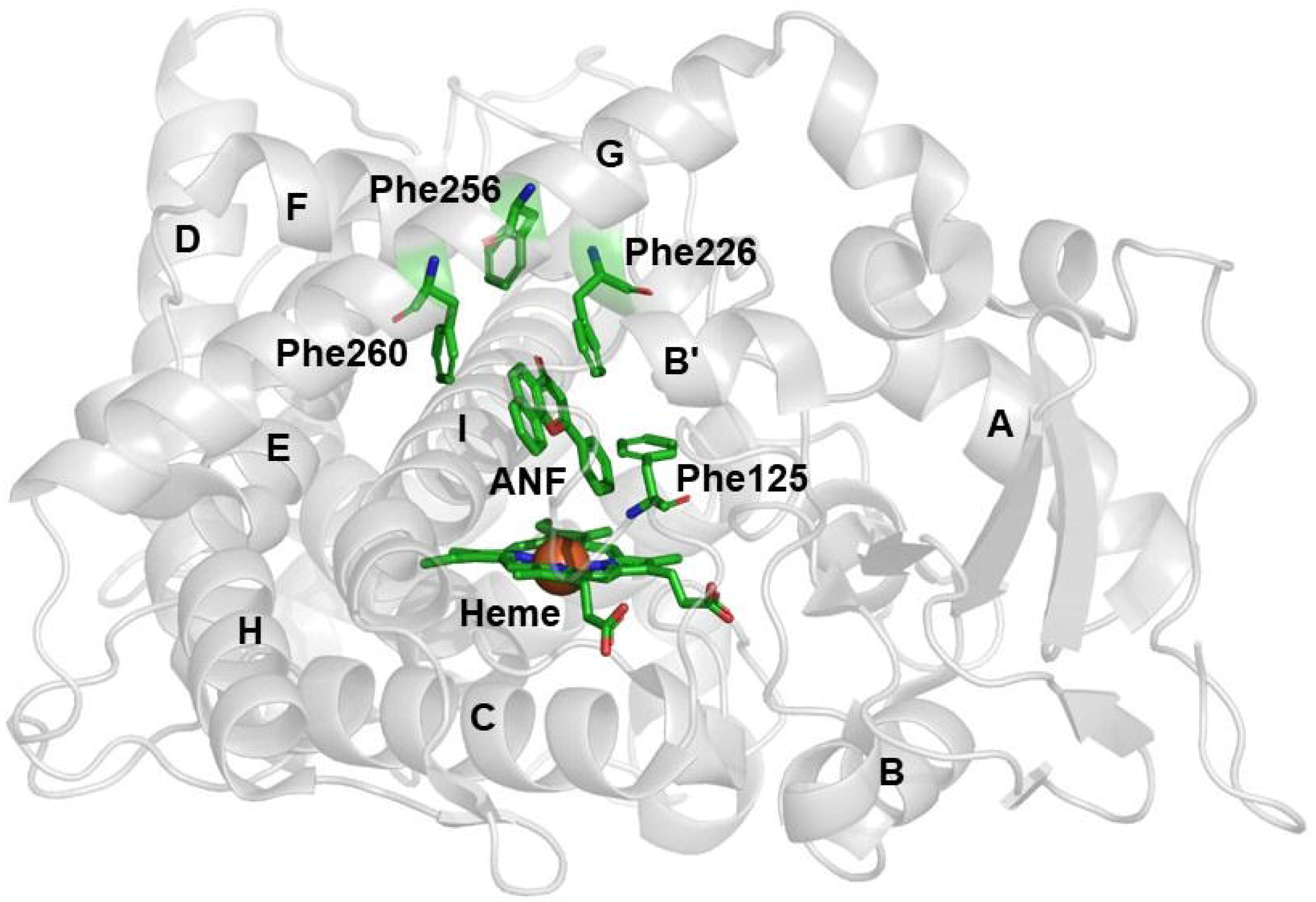
Illustratively showing the water channel in the wild type CYP1A2. (A) Left panel: the water channel is composed of BC loop, Right panel: the bottleneck radius of the water channel along 100-ns MD simulations; (B) Stereo-view of Ile386 location and water channel; (C) Stereo-view of Arg456 location and water channel.

Ile386 is located in the loop region between J and K helices (JK loop, **Figure 1**). Compared with other mutations (P42R, R431W, and R456H), I386F is not far away from the active pocket. This mutation is at the edge of the active pocket but has no interaction with the substrate. As shown in **Figure 5B**, Ile386 together with Thr124 forms the bottleneck of the water channel. The substitution from isoleucine to phenylalanine increases the sidechain length and forms a pi-pi stacking interaction with Phe125, which significantly reduces the radius of the bottleneck of the water channel.

Arg456 is located on the loop region between J and L helices (JL loop), residing below the heme plane (**Figure 1**). This residue forms two significant hydrogen bonds with Arg108 and Leu123 respectively, shaping the BC loop to form a stable water channel (**Figure 5C**). The substitution from arginine to histidine breaks the hydrogen bonds mentioned above, resulting in the instability of BC loop. The stability of the BC loop is a prior condition for a stable water channel. Thus, it is foreseeable that R456H can reduce the traversability of the substrate channel by increasing the instability of the BC loop.

To further confirm these points, we also calculated the bottleneck radius of the water channel along 100-ns MD simulations. In the current study, the water channel is considered open if the bottleneck radius is more than 1.5 Å. As a result, the water channel is open in most cases in the wild type CYP1A2 (**Figure 5A**). As expected, it becomes closed in most cases in both mutant structures, indicating that I386F and R456H mutations have negative impacts on the water channel.

### 4. R431W enhances the pi-pi stacking near the substrate channel

The substrate channel of the wild type CYP1A2 is composed of J helix, the loop region between J and K helices (JK loop), and the C-terminal loop region (**Figure 1**). As known, the loop regions are flexible in the solvent. In the wild type CYP1A2, multiple pi-pi stacking interactions are detected, from the protein surface to the active pocket, to position the flexible loop region around the substrate channel (**Figure 6A** and **Figure 2**). Thus, the conformations of the aromatic sidechains have a significant impact on the substrate channel.

**Figure 6.**
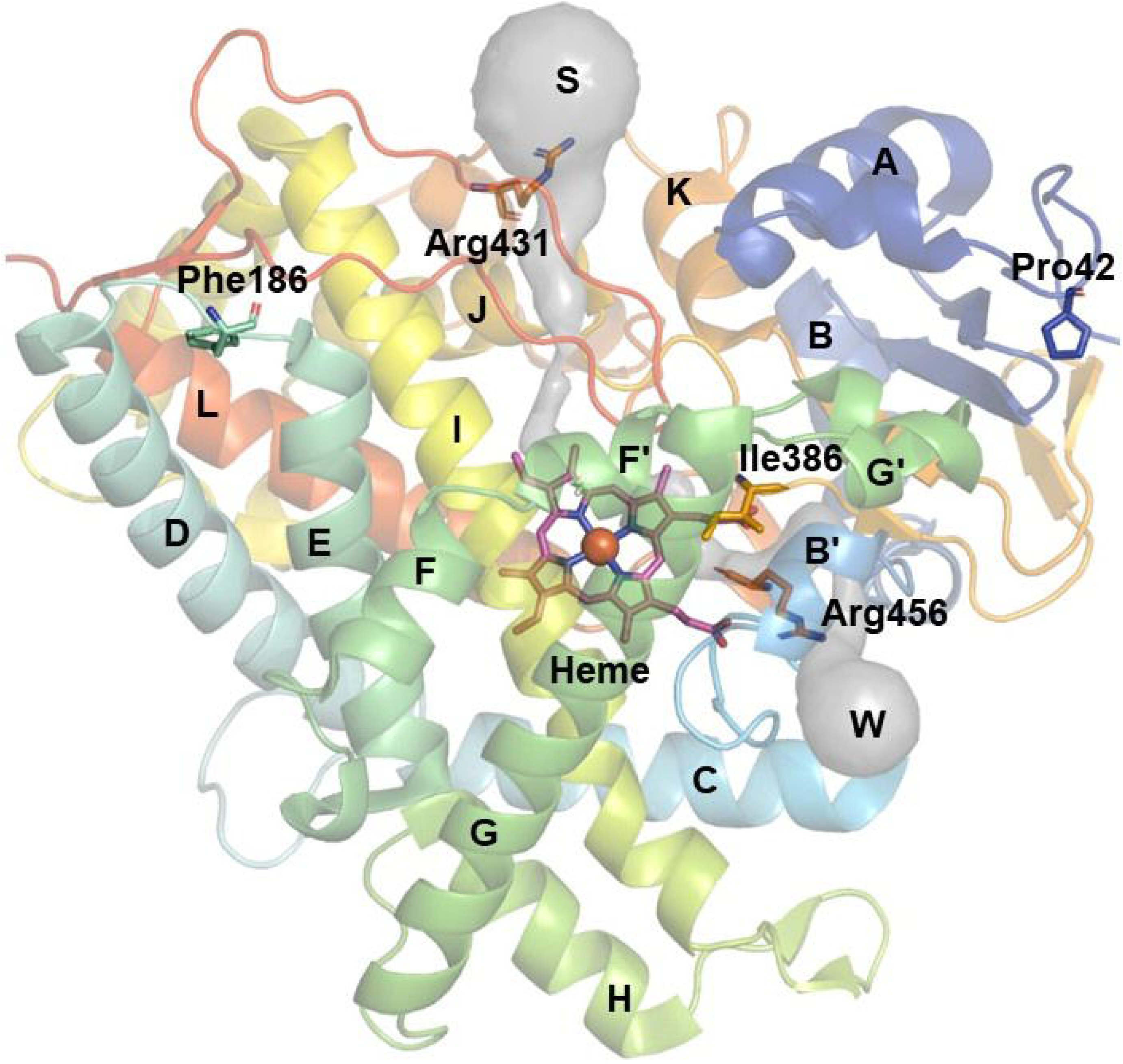
Illustratively showing the substrate channel in the wild type CYP1A2. (A) Multiple pi-pi stacking interactions around the substrate channel. (B) The bottleneck radius of the substrate channel along 100-ns MD simulations.

R431W is also a peripheral mutation located on K’ helix on the protein surface, 24.25 Å away from the heme iron (**Figure 1**). In the wild type CYP1A2, the sidechain of Arg431 has no influence on the multiple pi-pi stacking interactions mentioned above. In the R431W mutant structure, the substitution from arginine to tryptophan changes the aliphatic sidechain to aromatic, enhancing the multiple pi-pi stacking interactions. Due to locating on the protein surface, the aromatic sidechain of Trp431 is flexible in the solvent, resulting in the sidechain rearrangement of the aromatic residues via the multiple pi-pi stacking interactions.

To detect the impact of the sidechain rearrangement of the aromatic residues on the substrate channel, we calculated the bottleneck radius of the substrate channel along 100-ns MD simulations. In the current study, the substrate channel is considered open if the bottleneck radius is more than 4 Å. As a result, the substrate channel is open in most cases in the wild type CYP1A2 (**Figure 6B**). However, the substrate channel is closed in most cases in both F186L and R431W mutant structures (**Figure 6B**). This observation indicates that the R431W-induced sidechain arrangement of the aromatic residues reduces the open rate of the substrate channel, lowering the metabolic reactions of substrates.

## Discussion

CYP proteins are well-known for their single nucleotide polymorphisms or mutations. As estimated, the polymorphic variations or mutations of CYP proteins have a significant impact on more than 20% of clinical medicines (Chen et al., 2011; Wang et al., 2009). Because these mutations are capable of causing individual or population distinctions in the tolerance to clinical medicines. From a structural insight, these mutations that have a significant impact on the substrate metabolism can be classified into two types: central and peripheral mutation. The former is usually located in the active pocket of CYP proteins, having a crucial impact on the substrate binding. For instance, N404Y in human CYP2J2 decreases the binding affinity of substrates via breaking the hydrogen bonding interaction between Leu378 and the substrate (Cong et al., 2013). The latter is usually far away from the active pocket of CYP proteins, functioning to control the channels nearby. In human CYP1A2, F186L, I386F, R431W, and R456H are all peripheral mutations, having significant influences on the substrate or water channel. Different from these mutations, P42R is also far away from the active pocket but is adverse to membrane binding of CYP1A2.

In addition to locating away from the active pocket, another common feature for these mutations is involving in aromatic residues. Aromatic residues can form pi-pi stacking interactions, playing essential roles in recognizing and positioning substrates and forming access channels from the surface to the core region (Ping et al., 2012). Besides, aromatic residues also function to mediate cooperative binding properties for CYP proteins. For instance, Phe478 in human CYP2E1 is a key factor for multiple substrate binding, acting as a mediator for substrate communications via pi-pi stacking interactions (Li et al., 2011; Liu et al., 2013). The pi-pi stacking formed by Phe478 drives the substrates away from the catalytic center, going against the metabolic reactions. In human CYP3A4, Phe304 is a key factor for multiple substrate binding. This residue can enhance the pi-pi stacking interactions between substrates, which is propitious to substrate metabolisms (Fa et al., 2015).

## Conclusion

In conclusion, we constructed a membrane-binding model for full-length CYP1A2 and studied the dynamic behaviors of peripheral mutations of CYP1A2 (P42R, I386F, R431W, and R456H) on lipid membranes. Based on the computational simulations and free energy calculations, it is found that these peripheral mutations have no influence on the substrate binding. Instead, they can induce the access channel for water or substrates to adopt a closed state. These findings also provide a shred of evidence that pi-pi stacking interactions are essential for both substrate binding and channel opening in CYP proteins.

## Compliance with Ethical Standards

### Research involving human participants and/or animals

The article does not contain any studies with human participants or animals performed by any of the authors.

## Abbreviations

CYP: Cytochrome P450
SNP: single nucleotide polymorphisms
ANF: α-nappthoflavone
POPC: dioleoyl-sn-glycero-3-phosphoethanolamine
RMS: root-mean-square
MD: molecular dynamics
PME: particle mesh Ewald
MM-PB/SA: molecular mechanics Poisson-Boltzmann surface area
MM-GB/SA: molecular mechanics Generalized-Born surface area
PB: Poisson-Boltzmann
GB: Generalized Born
SASA: molecular solvent accessible surface areas
SAXS: small-angle X-ray scattering

